# Neuronal and oligodendroglial but not astroglial tau translates to in vivo tau-PET signals in primary tauopathies

**DOI:** 10.1101/2024.05.04.592508

**Authors:** Luna Slemann, Johannes Gnörich, Selina Hummel, Laura M. Bartos, Carolin Klaus, Agnes Kling, Julia Kusche-Palenga, Sebastian T. Kunte, Lea H. Kunze, Amelie L. Englert, Yunlei Li, Letizia Vogler, Sabrina Katzdobler, Carla Palleis, Alexander Bernhardt, Alexander Jäck, Andreas Zwergal, Franziska Hopfner, Sebastian Römer, Gloria Biechele, Sophia Stöcklein, Gerard Bischof, Thilo van Eimeren, Alexander Drzezga, Osama Sabri, Henryk Barthel, Gesine Respondek, Timo Grimmer, Johannes Levin, Jochen Herms, Lars Paeger, Marie Willroider, Leonie Beyer, Günter U. Höglinger, Sigrun Roeber, Nicolai Franzmeier, Matthias Brendel

**Author notes:** **Corresponding author:** Prof. Dr. Matthias Brendel, MHBA, Department of Nuclear Medicine, University of Munich Marchioninstraße 15, 81377 Munich, Germany. shared first authors (Luna Slemann, Johannes Gnörich, Selina Hummel, Laura M. Bartos). shared senior authors (Günter U. Höglinger, Sigrun Roeber, Nicolai Franzmeier, Matthias Brendel).

## Abstract

Tau-PET receives growing interest as an imaging biomarker for the 4-repeat tauopathy progressive supranuclear palsy (PSP). However, the translation of in vitro 4R-tau binding to in vivo tau-PET signals is still unclear. Therefore, we conducted a longitudinal [^18^F]PI-2620 PET/MRI study in a 4-repeat-tau mouse model (PS19) and found elevated [^18^F]PI-2620 PET signal in the presence of high neuronal tau. Cell sorting after radiotracer injection in vivo revealed higher tracer uptake in single neurons compared to astrocytes of PS19 mice. Regional [^18^F]PI-2620 tau-PET signals during lifetime correlated with abundance of fibrillary tau in subsequent autopsy samples of PSP patients and disease controls. In autoradiography, tau-positive neurons and oligodendrocytes with high AT8 density but not tau-positive astrocytes were the driver of [^18^F]PI-2620 autoradiography signals in PSP. In summary, neuronal and oligodendroglial tau constitutes the dominant source of tau-PET radiotracer binding in 4-repeat-tauopathies, yielding the capacity to translate to an in vivo signal.

## Introduction

Progressive supranuclear palsy (PSP) is a rare neurodegenerative disease caused by aggregation of tau protein with 4 microtubule-binding repeats (1), leading to death within 7-8 years after onset, on average (2). In the past, diagnosis was usually made 3-4 years after first clinical symptoms, when patients had already severe functional disabilities. Autopsy controlled data revealed that clinical diagnosis of PSP by most recent MDS-PSP criteria has only limited sensitivity in early stages at moderate specificity (3). Currently, no approved causal therapy exists and preventive and symptomatic treatment options are very limited. Since tau-targeting therapies are entering clinical development, it is now important to diagnose patients at an early stage of PSP to attenuate tau accumulation and therefore symptom progression. Furthermore, it is important to assure a high level of specificity for 4-repeat tau for inclusion in tau-targeting treatment trials to assure adequate statistical power and to avoid unnecessary side effects in clinically overlapping syndromes without 4-repeat tau aggregation. In this regard, cohort studies using the second generation tau-PET radiotracers [^18^F]PI-2620 and [^18^F]PM-PBB3 showed differentiation of patients with PSP and CBS from healthy and disease controls (4–6). However, head-to-head autoradiographic studies with different radiotracers by independent groups have been inconsistent, indicating the presence (7, 8) or absent (9) binding of radiolabeled PI-2620 binding to 4R-tauopathy tissue sections. Furthermore, potential off-target sources still need to be considered for second generation radiotracers, including neuromelanin and hemorrhagic lesions for [^18^F]PI-2620 (9), and in addition, β-amyloid for [^18^F]PM-PBB3 (10). Nevertheless, competitive assays (8) and molecular docking studies (11) confirmed the initially reported affinity of PI-2620 to 4R tau (12).

Hence, we aimed to investigate the translation of in vitro [^18^F]PI-2620 4R tau binding to in vivo PET signals. We tested longitudinal 4R tau monitoring in tau transgenic mice and pinpointed the cellular source of tracer signals by cell sorting after radiotracer injection in the living organism. Human in vivo PET signals in patients with definite PSP and disease controls were correlated with tau abundance and autoradiography signals in autopsy samples. Cellular and substructure impact to [^18^F]PI-2620 signal contributions were examined in depth using a PSP autoradiography sample with limited co-pathology. Finally, we exploited the combined acquired knowledge to create an optimized target region for detection of cortical tau pathology in patients with 4R-tauopathies.

## Results

### [^18^F]PI-2620 tau-PET monitoring reveals age-dependent increases of tracer binding in AT8 positive brain regions of the PS19 4-repeat tau mouse model

First, we investigated if [^18^F]PI-2620 tau-PET has sufficient sensitivity to detect an in vivo tracer signal in PS19 mice, which accumulate 4R tau pathology, over wild-type (WT) mice (**Fig. 1A**). Longitudinal tau-PET from 6 to 12 months of age showed an increasing PET signal and a significant genotype x age interaction effect in the entorhinal cortex (F(_3,40_) = 6.33, p = 0.0013) and the brainstem (F(_3,40_) = 9.09, p = 0.0001) of PS19 mice when compared to WT mice (**Fig. 1B-D**). The cerebellum did not qualify as a suitable target region in mice due to strong spill-over of adjacent skull relative to the brain (**Fig. 1B**). Late stage PS19 mice had a strongly elevated [^18^F]PI-2620 tau-PET signal in the entorhinal cortex (+19%, Cohen’s d = 3.26, p = 0.0003) and the brainstem (+21%, Cohen’s d = 2.47, p = 0.0009) compared to WT mice. Power calculation with standardized sample sizes of n=12 mice per genotype indicated an age of 9.7 months for earliest detection of signal alterations in cohorts of PS19 mice over WT at a predefined power of 0.8 (**Supplemental Fig. 2A,C**). Regional AT8 staining correlated with regional tau-PET signal enhancement at the final time-point (R = 0.921, p = 0.0032; **Supplemental Fig. 3)**. Volumetric 3T MRI analysis showed a decrease of hindbrain volumes in late stage PS19 mice when compared to WT mice (-7%, Cohen’s d = 1.98, p = 0.0062) and a weaker genotype x age interaction effect (F(_3,59_) = 3.93, p = 0.013) compared to tau-PET (**Fig. 1E,F**). Earliest detection of hindbrain volume differences was estimated at 11.7 months of age (**Supplemental Fig. 2B,C**). PS19 mice with high tau-PET signals in the brainstem were characterized by strong volume loss in the brainstem, whereas this association was not present in WT mice (**Fig. 1G**).

**Figure 1.**
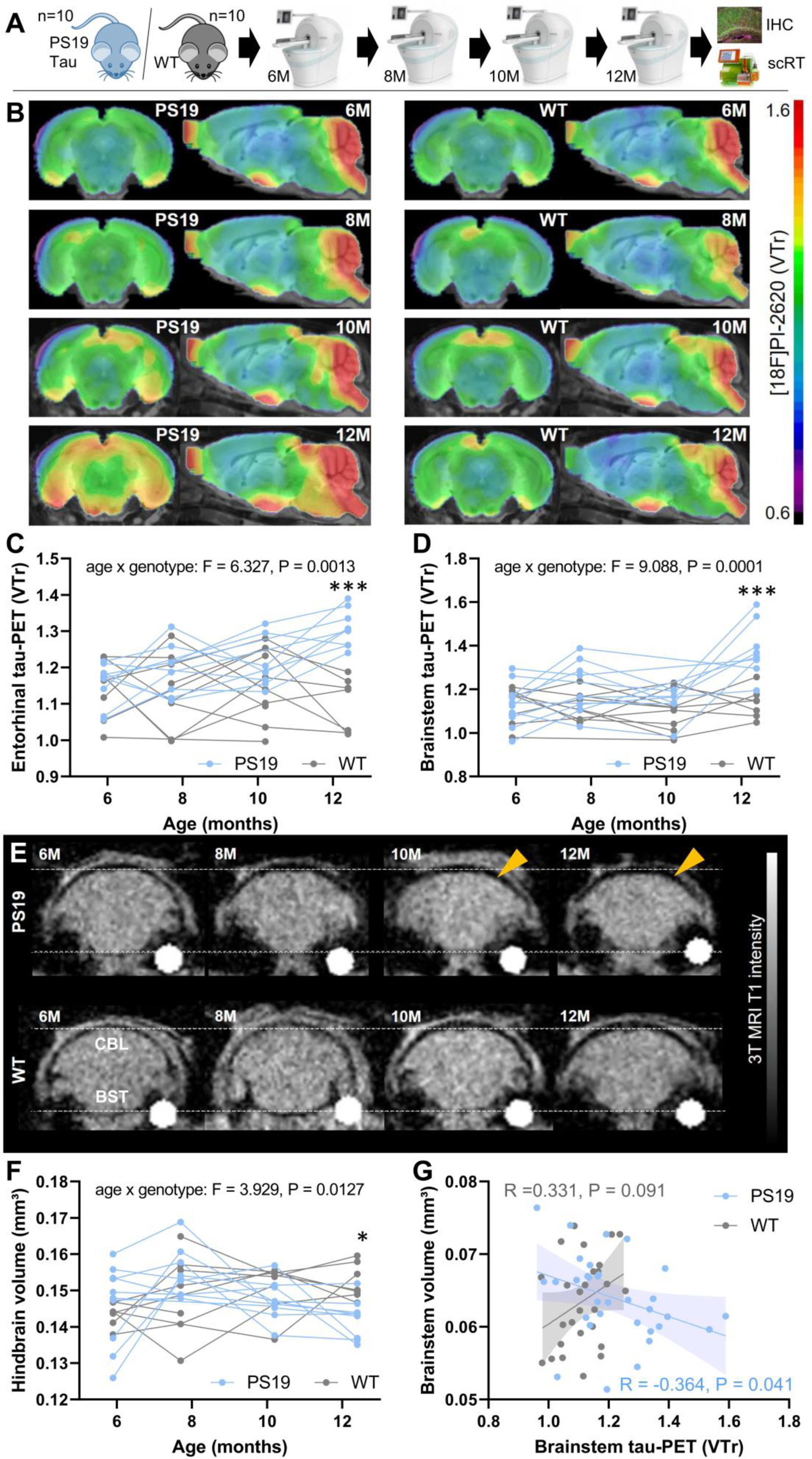
Monitoring of tau pathology and atrophy in PS19 and wild-type mice using [^18^F]PI-2620 PET/MRI. (**A**) Experimental workflow of serial PET/MRI imaging sessions and terminal immunohistochemistry (IHC) and single cell Radiotracing (scRT) in PS19 and wild-type (WT) mice. (**B**) Coronal and axial group average [^18^F]PI-2620 PET images upon a MRI template show monitoring of radiotracer binding (volume of distribution ratios, VT; striatal reference) with pronounced temporal and brainstem patterns in aged PS19 mice (n = 8-10) compared to WT mice (n = 8-10). (**C,D**) Mixed linear models of entorhinal and brainstem [^18^F]PI-2620 PET binding indicate a significant age x genotype effect and elevated tau-PET signals of PS19 compared to WT mice at 12 months of age. (**E**) Examples of serial MRI atrophy patterns in an individual PS19 mouse compared to a WT mouse. Coronal slices are shown with indication of cerebellum (CBL) and brainstem (BST) regions. Orange arrows highlight atrophy in PS19 mice. (**F**) Mixed linear model of hindbrain volume indicates a significant age x genotype effect and decreased hindbrain volume of PS19 compared to WT mice at 12 months of age. (**G**) Association between tau-PET binding and brain volume in the brainstem of PS19 and WT mice across all investigated time-points shows higher atrophy in presence of high tau-PET signals specifically in PS19 mice.

### Immunohistochemistry shows dominance of neuronal over astrocytic tau in PS19 mice

Next, we assessed the detailed regional sources of tau pathology in the brain of PS19 mice using immunohistochemistry. In line with previous work, high abundance of AT8-positive tau pathology was observed in hippocampus (+20.1%, p = 0.0004), cortex (+7.0%, p = 0.0025) and brainstem (+5.0%, p = 0.0321) of PS19 mice at 12 months, compared to age-matched WT mice (**Fig. 2A,B**) (13). Contrary, the cerebellum indicated no significant elevation of stained AT8-positive area (1.3%, p = 0.552). Similarly, stronger GFAP reactivity of PS19 mice over WT as a surrogate of reactive astrogliosis was observed in the frontal cortex (+9.7%, p = 0.025) and the hippocampus (+9.2%, p = 0.050; **Fig. 2A,C**) (14).

**Figure 2.**
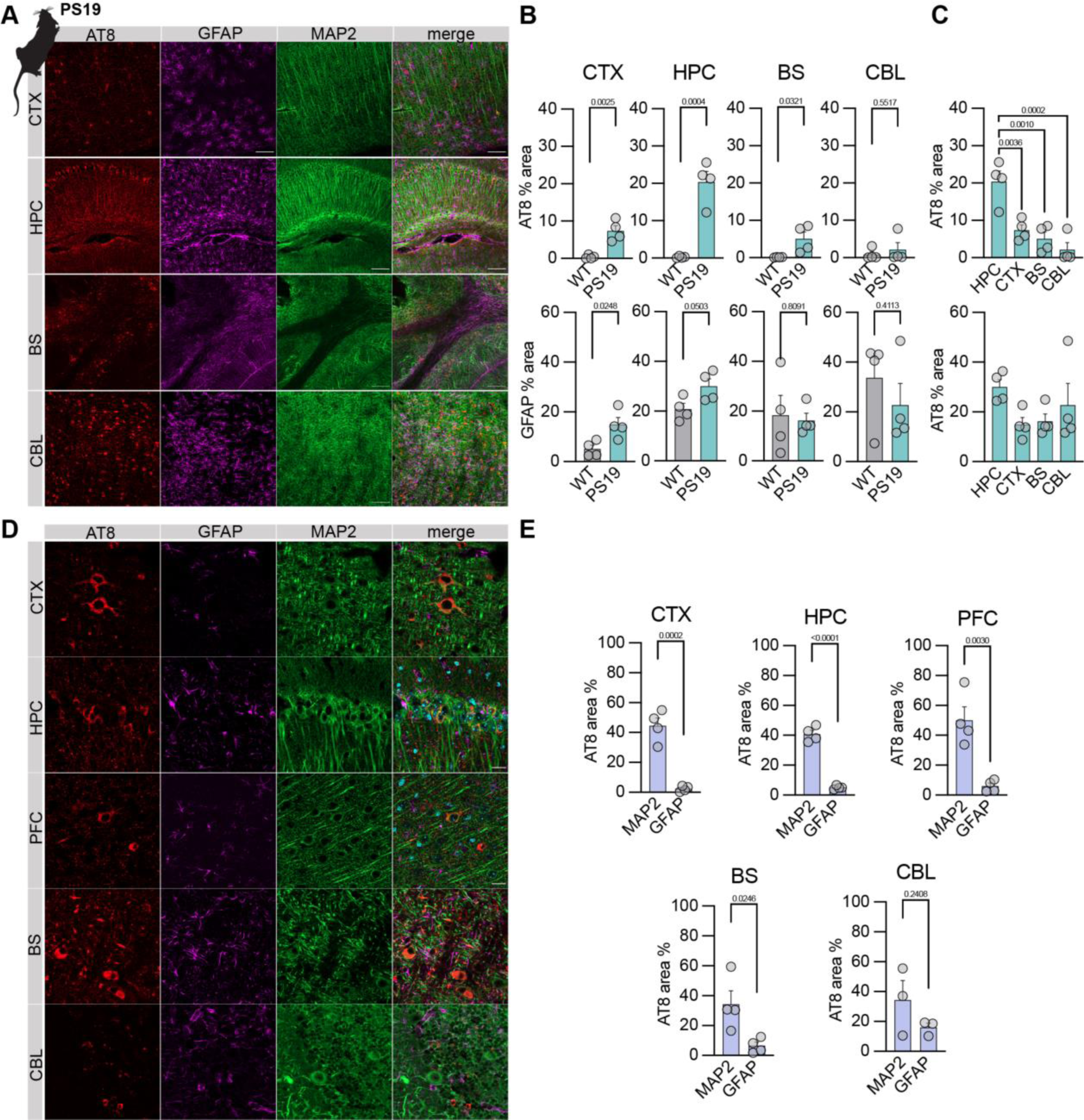
Immunohistochemistry assessment of tau pathology, reactive astrocytes and neurons in PS19 mice. (**A**) Overview sections of PS19 mice show pTau (AT8, red), astrocytes (GFAP, purple), and neurons (MAP2, green) as well as merged images taken from cortex (CTX), hippocampus (HPC), cerebellum (CBL) and brainstem (BS). Scale bar = µm. (**B**) Quantitative comparison of AT8 occupancy in target regions between wild-type (WT) and PS19 mice as well as comparison of PS19 AT8 occupancy across target regions. (**C**) Quantitative comparison of GFAP occupancy in target regions between wild-type (WT) and PS19 mice as well as comparison of PS19 GFAP occupancy across target regions. (**D**) Zoomed sections of PS19 mice show pTau (AT8, red), astrocytes (GFAP, purple), and neurons (MAP2, green) as well as merged images taken from cortex (CTX), hippocampus (HPC), cerebellum (CBL) and brainstem (BS). Scale bar = µm. (**E**) Quantitative comparison of AT8 occupancy between neurons (MAP2-positive) and astrocytes (GFAP-positive) in target regions of PS19 mice. CTX =cortex, HIP = hippocampus, PFC = prefrontal cortex, BS = brainstem, CBL = cerebellum.

Co-labeling of AT8 with GFAP (astroglia) and MAP2 to mark neuronal soma including somatodendritic structures revealed that tau aggregation predominantly occurred in neurons (somata and putatively postsynaptic compartments) but to a significantly lower degree in astrocytes (2-18 fold, all target regions p < 0.05, **Fig. 2X**). Collectively, these data reflect signals obtained in PS19 mice subjected to tau PET imaging and depicts the predominant neuronal origin of tau pathology.

### Increased neuronal [^18^F]PI-2620 uptake of late-stage PS19 mice translates to an in vivo PET signal

To determine the cellular source of [^18^F]PI-2620 binding in PS19 mice, we performed scRadiotracing in a subset of five transgenic and five WT mice immediately after the final tau-PET session (**Fig. 3A**). Isolation of neurons and astrocytes was performed using MACS and radioactivity was measured in enriched cell pellets. Strikingly, we observed a 1.89-fold higher [^18^F]PI-2620 uptake per single neuron of PS19 mice compared to single neurons of WT mice (p = 0.028; **Fig. 3B**). Contrary, [^18^F]PI-2620 uptake per single astrocytes was not different between PS19 and WT mice (p = 0.439; **Fig. 3C**). In PS19 mice, neuronal tracer uptake was 27-fold higher when compared to astrocytes (p = 0.023), while WT mice also indicated higher [^18^F]PI-2620 uptake of neurons when compared to astrocytes (5-fold, p = 0.009). Individual tau-PET z-score maps of PS19 mice matched the magnitude of single cell [^18^F]PI-2620 uptake in neurons (**Fig. 3D**). Furthermore, late static tau-PET quantification in a predefined hindbrain region was correlated with individual neuronal tracer uptake of the whole sample (R = 0.727, p = 0.017) and in PS19 mice (R = 0.919, p = 0.027; **Fig. 3E**), whereas no significant associations were found between tau-PET and individual astrocytic tracer uptake (**Fig. 3F**). To allow more regional flexibility, we correlated the individual neuronal and astrocytic tracer uptake with voxel-wise tau-PET using SPM. Neuronal [^18^F]PI-2620 uptake was correlated with regional PET signals in brainstem, midbrain and entorhinal cortex (**Fig. 3G**) whereas astrocytic [^18^F]PI-2620 uptake did not show any significant associations with regional tau-PET signals (**Fig. 3H**). Flow cytometry confirmed sufficient cellular yield (**Fig. 3I**) and purity (**Fig. 3J**) of neurons and astrocytes. Thus, we questioned if the magnitude of cell specific tracer uptake in individual PS19 animals corresponds to the signal alterations observed in PET. Considering 71*10e^6^ neurons and 21*10e^6^ astrocytes in the rodent brain, we found that cellular radioactivity (7.7-36.5 kBq per animal) explained the specific tau-PET signal enhancement (16.4-37.2 kBq per animal; p=0.880), also resulting in closely matching summations of cellular and PET related radioactivity measures across the five PS19 mice studied (**Fig. 3K**).

**Figure 3.**
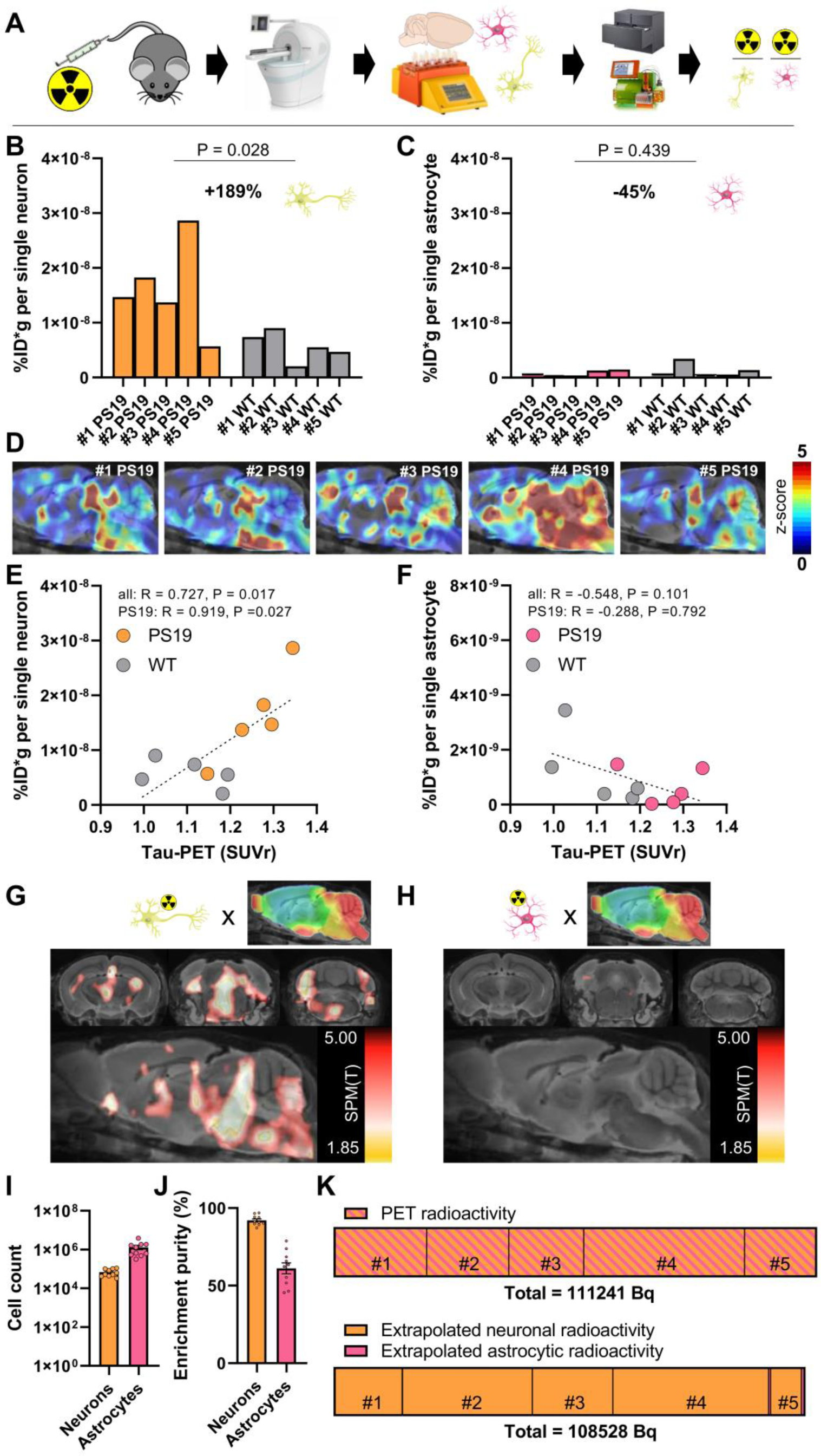
Cell sorting after radiotracer injection identifies neurons as the predominant origin of [^18^F]PI-2620 tau-PET signals. (**A**) Experimental workflow of PET/MRI imaging with subsequent cell sorting of neurons and astrocytes prior to determination of radioactivity per isolated cell by gamma emission measures and flow cytometry. (**B,C**) Radioactivity per isolated neuron and astrocyte in comparison of PS19 (n=5) and wild-type (WT, n=5) mice. Each bar represents an individual animal. (**D**) Sagittal sections upon a MRI template show z-score images (vs. WT) of all investigated PS19 mice (n=5). Each image represents an individual animal. (**E,F**) Quantitative correlation between brainstem tau-PET signals and radioactivity per single neuron or astrocyte. Person’s coefficients of correlation are provided for the combined data of PS19 and WT mice (regression line with 95% confidence interval) as well as for the subset of PS19 mice. (**G,H**) Data driven voxel-wise correlation between radioactivity per single neuron or astrocyte and [^18^F]PI-2620 tau-PET images using statistical parametric mapping of the combined sample of PS19 and WT mice. Radiotracer uptake per neuron correlated with the distribution pattern of tau pathology in PS19 mice, whereas radiotracer uptake per astrocyte did not show a correlation with tau-PET patterns. (**I,J**) Cell count and purity of isolated cells (neurons and astrocytes) as benchmark indices of the cell sorting procedure. (**K**) Quantitative PET radioactivity increases in PS19 mice (n=5) compared to WT mice match radioactivity as determined by isolated cells. Radioactivity per single cell was extrapolated using established cell numbers of the mouse brain (71 × 10e^6^ neurons, 21 × 10e^6^ astrocytes).

### [^18^F]PI-2620 tau-PET signals correlate strongly with regional tau abundance in deceased patients with PSP and disease controls

Next, we examined if in vivo signals of the tau-PET tracer [^18^F]PI-2620 are determined by tau neuropathology. To this end, we investigated a small cohort of eight patients that underwent [^18^F]PI-2620 PET in vivo, with subsequent donation of their brains for autopsy. Six patients were classified as definite PSP and two patients were classified as TAR DNA-binding protein 43 (TDP-43)-positive frontotemporal lobar degeneration FTLD-TDP: one FTLD/MND-TDP; one FTLD-TDP related to a TANK-binding kinase 1 (TBK1) mutation (**Supplemental Table 1**). The globus pallidus showed higher visual and quantitative AT8 occupancy compared to the medial frontal gyrus in patients with definite PSP, which was well reflected by corresponding autoradiography signals (**Fig. 4A,B**). Tau-PET, autoradiography and AT8 immunohistochemistry indicated higher signals or occupancy in patients with definite PSP compared to disease controls (**Fig. 4A,B**). Noteworthy, the TBK1 mutation carrier showed mild AT8-positive tau co-pathology in the globus pallidus and in the putamen and moderate AT8-Co-pathology in basal nucleus Meynert at autopsy, which explained moderate elevation of the [^18^F]PI-2620 PET signal in these regions. Across modalities, there was a strong correlation between quantitative AT8 occupancy and autoradiography ratios (R = 0.943, p < 0.0001) even when challenged by combined consideration of both target regions (**Fig. 4C**). Furthermore, quantitative AT8 occupancy (R = 0.815, p = 0.0012) and autoradiography ratios (R = 0.886, p = 0.0001) showed high agreement with tau-PET signals in vivo, acquired 5-39 months before death (**Fig. 4C**).

**Figure 4.**
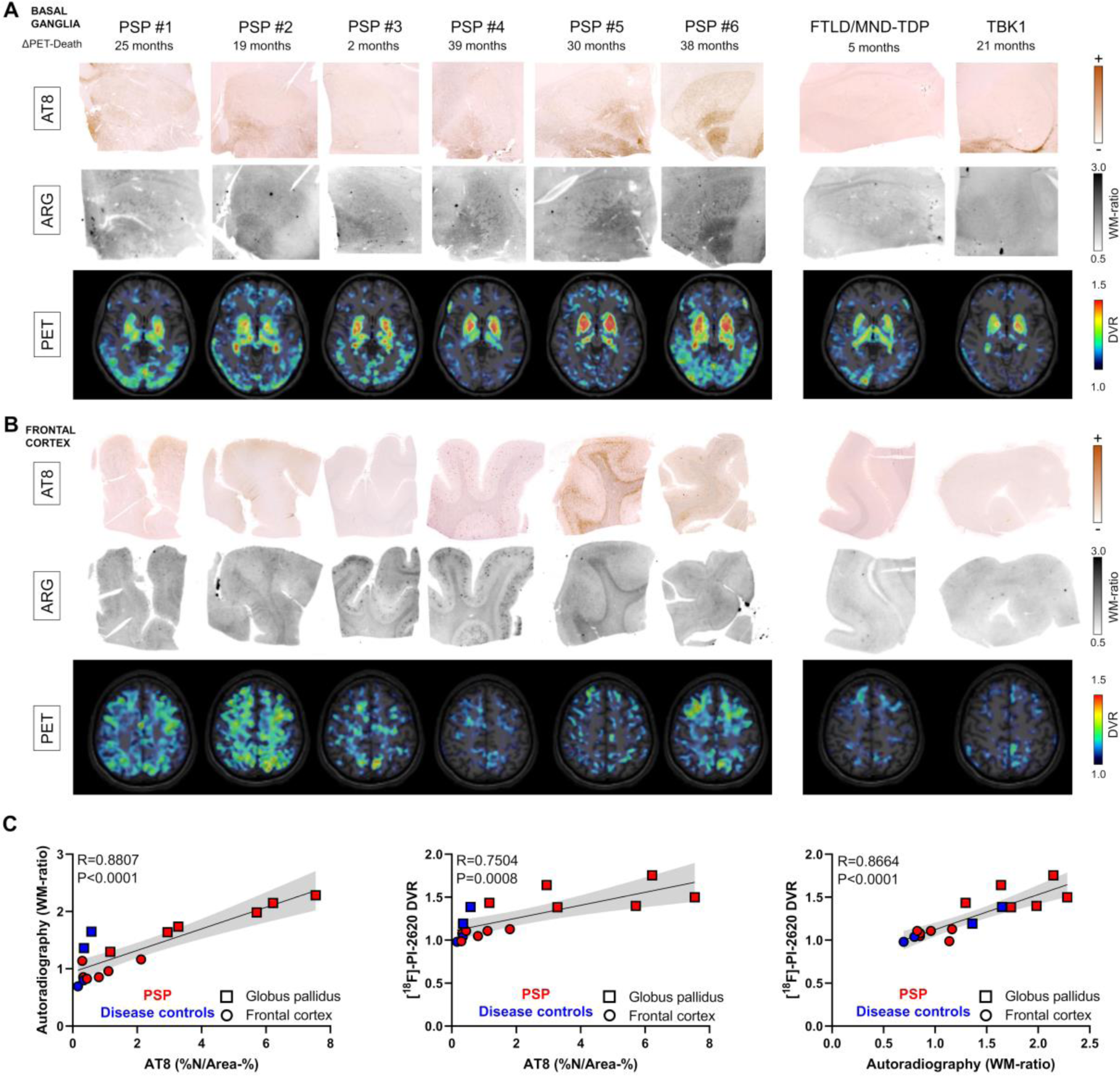
Correlation of in vivo PET signals with tau abundance and autoradiography signals in autopsy samples. (**A**) Basal ganglia AT8 immunohistochemistry together with [^18^F]PI-2620 autoradiography of adjacent sections and axial [^18^F]PI-2620 tau-PET basal ganglia sections prior to death. Images display all six investigated patients with definite PSP and two patients with TAR DNA-binding protein 43 (TDP-43)-positive frontotemporal lobar degeneration (FTLD-TDP): one FTLD/MND-TDP; one FTLD-TDP related to a TANK-binding kinase 1 (TBK1) mutation. (**B**) Frontal medial gyrus AT8 immunohistochemistry together with [^18^F]PI-2620 autoradiography of adjacent sections and axial [^18^F]PI-2620 tau-PET cortex sections prior to death. Images display six patients with definite PSP and two disease controls as indicated in (A). (**C**) Multimodal quantitative correlation between tau abundance, autoradiography signals and tau-PET signals. Regression lines (including 95% confidence intervals) were calculated for the combined analysis of globus pallidus and frontal medial gyrus samples derived from six patients with definite PSP, one patient with FTLD/MND-TDP and one patient with FTLD related to a TBK1 mutation. R indicates Pearson’s coefficient of correlation. %N/Area-% = AT8 occupancy of neurofibrillary tangles and coiled bodies.

### In vitro autoradiography confirms tau-positive neurons and oligodendrocytes as the major source of tau-tracer binding in PSP tissue

Next, we translated our murine cell-type findings to human tau-PET imaging and examined the detailed sources of tau-PET signals. A correlation analysis between the area of AT8-positive neurons/oligodendrocytes (NFT/CB) as well as the area of AT8-positive astrocytes (TA, including TF) and [^18^F]PI-2620 autoradiography signals was performed using autopsy tissue derived from 16 patients with PSP (**Supplemental Table 2**). This PSP sample was selected based on absence of α-synuclein, TDP-43 or FUS co-pathology and limited β-amyloid co-pathology to minimize confounding factors, as assessed in the frontal cortex.

The frontal cortex was used as primary brain region of interest due to low probability of off-target sources (15). Here, we found substantial visual agreement between AT8 immunohistochemistry and [^18^F]PI-2620 autoradiography signals in FFPE sections of patients with definite PSP (**Fig. 5A**). We tested the differential associations of neuronal/oligodendroglial (NFT/CB) and astrocytic (TA, including TF) tau abundance, as determined by AT8 immunohistochemistry (**Fig. 5B**), with [^18^F]PI-2620 autoradiography quantification in 129 predefined subfields of grey matter and white matter (drawn on autoradiography sections blind to the corresponding AT8 sections; **Supplemental Fig. 4**). NFT/CB tau abundance in PSP samples correlated stronger with autoradiography quantification (R = 0.487, p < 0.0001, **Fig. 5C**) compared to TA/TF tau abundance (R = 0.280, p = 0.0013, **Fig. 5D**). A regression analysis with NFT/CB and TA/TF tau abundance as predictors showed that only NFT/CB tau (β = 0.455,p < 0.0001) but not TA/TF tau (β = 0.068, p = 0.442) explained the [^18^F]PI-2620 autoradiography signal. We noticed that subfield tau abundance in individual PSP samples showed substantial correlation with autoradiography signals only above 0.2% mean occupied area of tau-positive NFT/CB (R = 0.623, p = 0.017, **Fig. 5E**), indicating the sensitivity threshold for translation into measurable signal. Contrary, individual PSP samples with high mean occupied area of tau-positive TA/TF did not show strong individual subfield correlations (**Fig. 5F**), again arguing for limited translation of TA/TF tau to autoradiography and PET signals.

**Figure 5.**
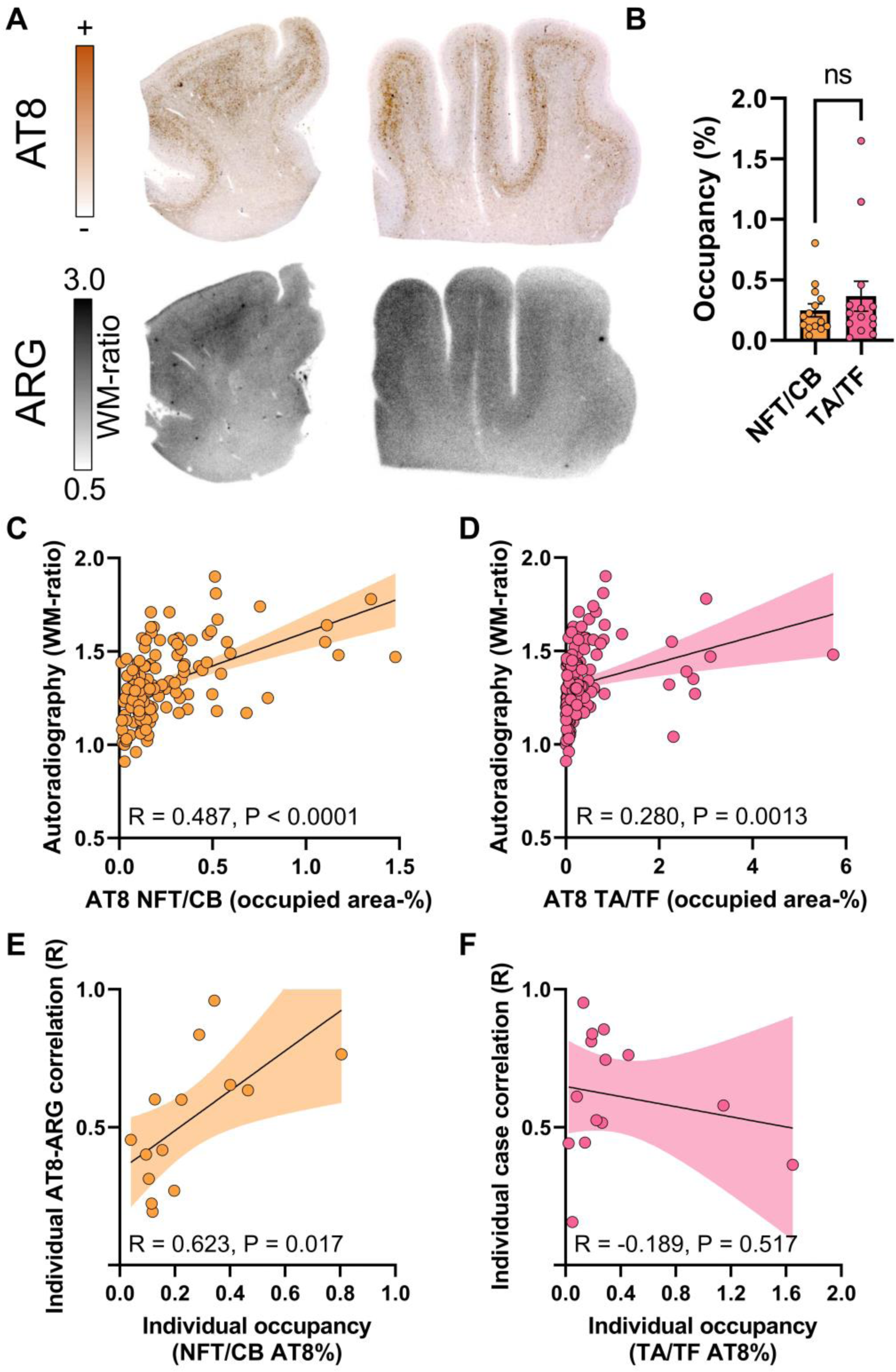
Cell type specific immunohistochemistry to autoradiography correlation in the frontal medial gyrus of PSP tissue samples. (**A**) AT8 immunohistochemistry together with [^18^F]PI-2620 autoradiography of adjacent frontal medial gyrus sections of two exemplary patients with definite PSP. ARG = autoradiography. WM = white matter (**B**) Comparison of neuronal/oligodendroglial (NFT/CB) and astrocytic (TA, including TF) AT8 occupancy in frontal medial gyrus sections of patients with definite PSP (n=14). (**C, D**) Quantitative correlation between tau abundance of NFT/CB and TA/TF with autoradiography signals in frontal medial gyrus subfields (including grey matter and white matter regions). Regression lines (including 95% confidence intervals) were calculated for 129 subfields derived from n=14 patients with definite PSP. R indicates Pearson’s coefficient of correlation. (**E, F**) Dependency of individual subfield correlation coefficients (AT8 x autoradiography) from mean AT8 occupancy of all subfields per patient as determined for NFT/CB and TA/TF. Regression lines (including 95% confidence intervals) were calculated n=14 patients with definite PSP. R indicates Pearson’s coefficient of correlation.

Despite substantial visual agreement in most PSP samples (**Fig. 6A**) and presence of predominant TA/TF tau aggregation compared to sparse NFT/CB tau aggregation (**Fig. 6B**), only NFT/CB tau occupancy showed a substantial correlation with [^18^F]PI-2620 autoradiography signals in 73 subfields of the globus pallidus (**Fig. 6C**, R = 0.467, p < 0.0001). Contrary, the quantitative agreement between TA/TF tau occupancy and [^18^F]PI-2620 autoradiography signals only reached borderline significance (**Fig. 6D**, R = 0.231, p = 0.0498). Again, a regression model including NFT/CB AT8 occupancy and TA/TF AT8 occupancy as predictors showed only a significant explanation of autoradiography signals by NFT/CB tau (β = 0.676, p < 0.0001), but not by TA/TF tau (β = -0.278, p = 0.081). Compared to the frontal cortex, AT8-positive areas in the basal ganglia were characterized by more heterogeneity of AT8 staining intensity, including high abundance of AT8 positive axons and occupancy of large neurons with high AT8 intensity in individual cases (see example in **Fig. 6E-G**). Notably, large AT8-positive neurons (NFT) were visually discernible in [^18^F]PI-2620 autoradiography images (**Fig. 6E**). We questioned the impact of AT8 density on the resulting autoradiography signal and observed a strong association between AT8-lesion density and autoradiography signals in individual patients (R = 0.800, p = 0.017, **Fig. 6F**), whereas the AT8-occupied area was no significant predictor of the autoradiography signal in such cases (**Fig. 6G**). Noteworthy, some PSP samples showed a visually detectable [^18^F]PI-2620 autoradiography signal in the putamen although AT8 occupancy was low, fitting to higher yet unexplained background [^18^F]PI-2620 signals in the putamen of healthy controls compared to cortical areas (4).

**Figure 6.**
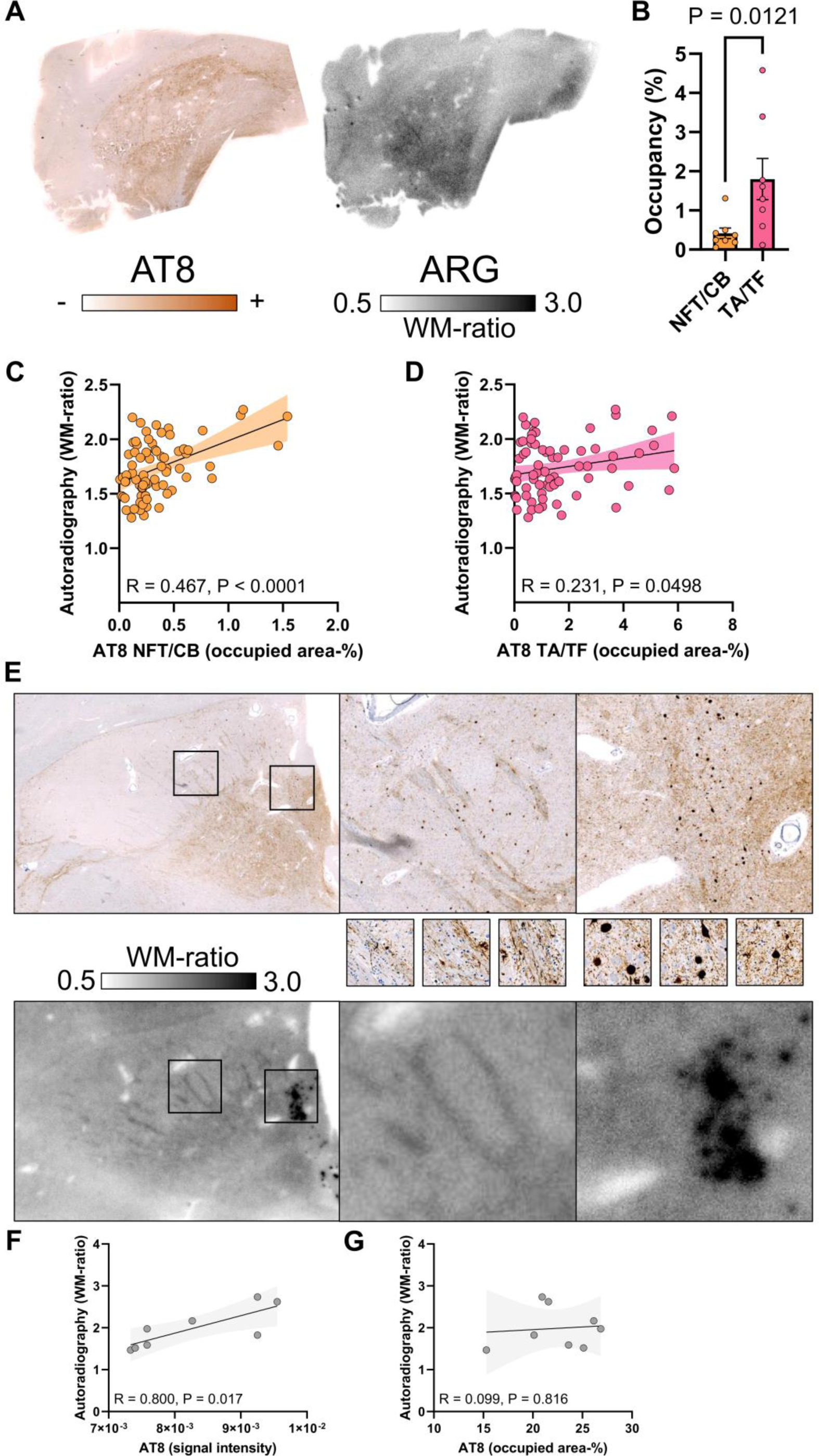
Cell- and substructure-specific immunohistochemistry to autoradiography correlation in the basal ganglia of PSP tissue samples. (**A**) AT8 immunohistochemistry together with [^18^F]PI-2620 autoradiography of the adjacent basal ganglia section of one exemplary patient with definite PSP. (**B**) Comparison of neuronal and oligodendroglial (NFT/CB) versus astrocytic (TA, including TF) AT8 occupancy in the globus pallidus of patients with definite PSP (n=8). (**C**) Quantitative correlation between NFT/CB tau abundance and autoradiography signals in basal ganglia subfields. Regression line (including 95% confidence intervals) was calculated for 73 subfields derived from n=8 patients with definite PSP. R indicates Pearson’s coefficient of correlation. (**D**) Quantitative correlation between TA/TF tau abundance and autoradiography signals in basal ganglia subfields. Regression line (including 95% confidence intervals) was calculated for 73 subfields derived from n=8 patients with definite PSP. R indicates Pearson’s coefficient of correlation. (**E-G**) Quantitative and visual correlation between AT8 intensity and AT8 occupancy with autoradiography signals in an individual PSP sample. Small images in (**E**) show zoom of faint AT8 intensity in white matter fibers and strong AT8 intensity in large neurons (NFTs). Note the strong autoradiography signal which is discernible as single spots in the same area. Regression lines (including 95% confidence intervals) were calculated for n=8 subfields of one patient with definite PSP. R indicates Pearson’s coefficient of correlation.

### High oligodendroglial density at the boundary of grey and white matter points to an improved cortical target region for tau-PET in PSP

Finally, we exploited cellular and regional findings of our study to further improve tau-PET diagnostic in 4R-tauopathies. Pronounced regional autoradiography signals were observed in white matter areas adjacent to the GM/WM boundary (**Fig. 7A**). In line with previous reports (16, 17), these areas were characterized by distinctly higher NFT/CB-to-TA/TF ratios when compared to cortical grey matter regions (4-fold, p < 0.0001, **Fig. 7B**). To test for the relevance of this observation in vivo, we analyzed seventeen [^18^F]PI-2620 tau-PET scans of patients with PSP-RS and high likelihood of underlying 4R-tauopathy and nine healthy controls with a MRI-based layer segmentation of grey matter and WM/GM boundary areas of the frontal cortex. Here, we found higher effect size for the comparison of tau-PET signals in the frontal cortex of patients with 4R-tauopathies against controls by use of the GM/WM boundary target region (Cohen’s d = 1.68) in contrast to the common grey matter target region (Cohen’s d = 1.37). Thus, focusing on oligodendroglia-rich regions with high AT8-positivity enhanced assessment and diagnostic accuracy of cortical tau burden by [^18^F]PI-2620 tau-PET in patients with 4R-tauopathies.

**Figure 7.**
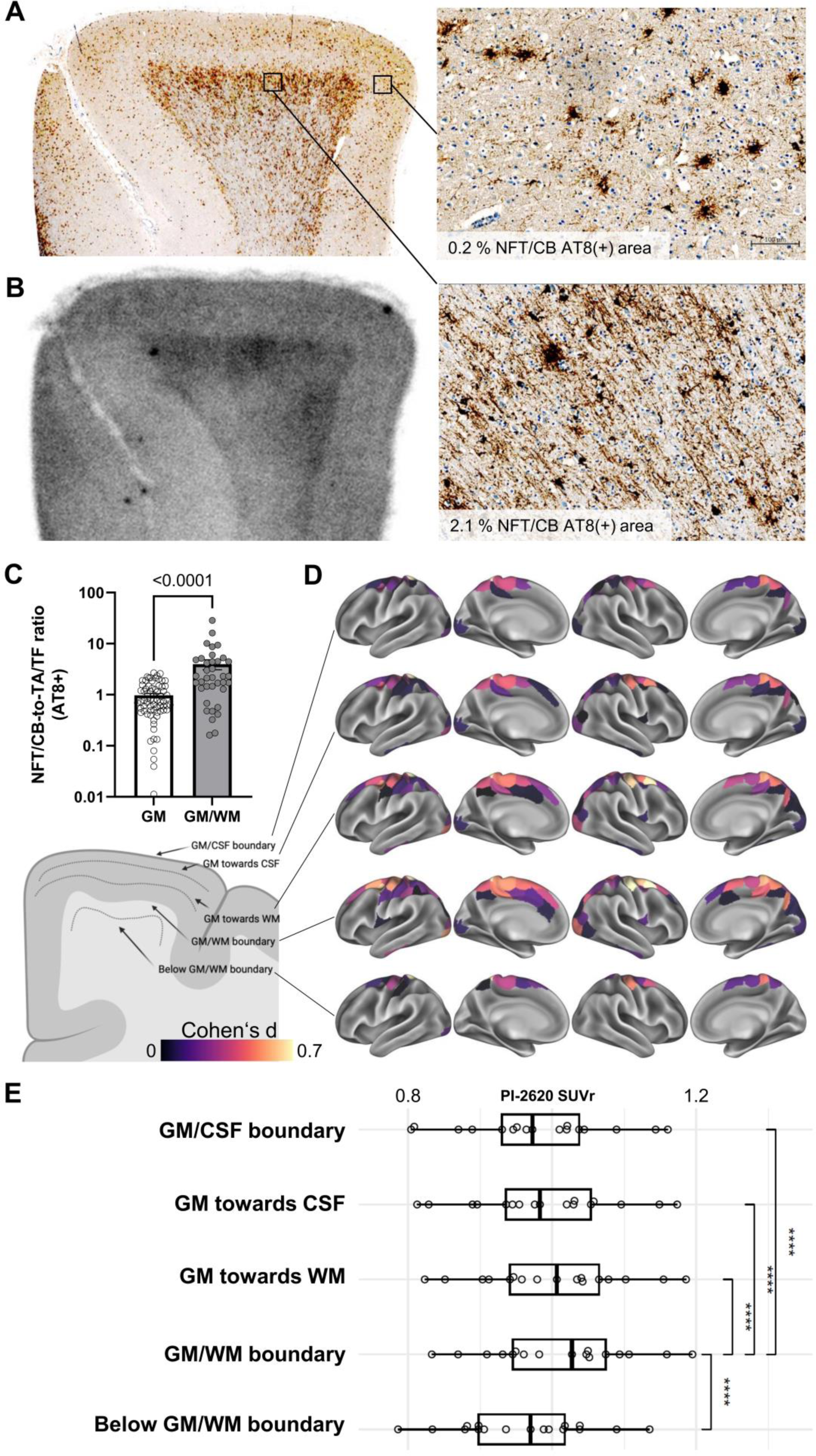
High oligodendroglial tau abundance in the GM/WM boundary facilitates definition of an optimized frontal lobe PSP target region. (**A,B**) AT8 immunohistochemistry together with [^18^F]PI-2620 autoradiography of the adjacent frontal medial gyrus section of one exemplary patient with definite PSP. Zoom images show high oligodendroglial tau in the boundary of grey matter and white matter, whereas cortical layers are characterized by low neuronal/oligodendroglial (NFT/CB) tau but abundant astrocytic (TA) tau inclusions. (**C**) Quantitative comparison of NFT/CB to TA/TF ratios of tau occupancy in the grey matter and the boundary of grey matter and white matter. Data derive from NFT/CB and TA/TF AT8 occupancy in frontal medial gyrus subfields of patients with definite PSP (n=14). (**D**) Schematic illustration of PET SUVR assessment from the GM/WM boundary to the GM/CSF boundary. Surface renderings illustrate standardized group differences (Cohen’s d) between 17 PSP-RS patients and 9 healthy controls for the 200 regions of the Schaefer cortical atlas. Group-differences are shown across 5 MRI-based cortical surface reconstructions applied to the PET data, systematically shifted from the GM/CSF boundary towards the GM/WM boundary and below. (**E**) Boxplots show corresponding PET SUVRs in the 17 PSP-RS patients.

## Discussion

In this translational study, we used a large spectrum of methodological approaches, including innovative scRadiotracing and cell type specific correlation of autoradiography signals, to disentangle discrepant findings of previous reports that investigated second generation tau-PET in 4R-tauopathies. As a major achievement, we were able to detect elevated radiotracer binding in isolated neurons after in vivo injection in mice. Furthermore, our data indicate that tau-PET signals in 4R-tauopathies are driven by neuronal and oligodendroglial tau aggregation whereas faint tau-positive structures of astrocytes and tau fragments are not capable to translate radiotracer binding into in vivo signaling. Finally, we provide first [^18^F]PI-2620 PET to autopsy correlation and show that cortical tau-PET signals deserve optimized target regions at the boundary between grey and white matter.

The overarching research question of this work addressed the validity of second generation tau-PET signals in 4R-tauopathies. A recent blocking study found that [^3^H]PI-2620 but not [^3^H]MK-6240 or [^3^H]RO-948 indicated high specific binding in frontal cortex of deceased patients with PSP and CBD (7). Contrary, another recent autoradiography head-to-head comparison did not find a significant extent of [^18^F]AV-1451, [^18^F]MK-6240 or [^18^F]PI-2620 binding to non-AD tauopathies (9). Such discrepancies were already reported earlier for [^18^F]AV-1451, showing positive (18) and negative (19) autoradiography signals in 4R-tauopathy brain sections. However, in addition, in vitro saturation assays and competitive binding assays (8) as well as molecular docking (11) provided further evidence of PI-2620 binding to 4R-tau. Therefore, we have carried out a battery of experiments in order to test the translation of PI-2620 binding to in vivo PET signals.

First, we were able to show that second generation tau-PET with [^18^F]PI-2620, a radiotracer that indicated affinity to 3- and 4-repeat tau, has sufficient sensitivity to detect accumulating tau pathology in transgenic PS19 mice. Tau-PET indicated earlier sensitivity for pathological alterations when compared to 3T structural MRI based atrophy measures, but we note that both modalities could still be improved in terms of resolution, i.e. availability of ultrahigh field MRI. In terms of preclinical tau-PET imaging, [^18^F]PM-PBB3 also showed elevated tau-PET signals in the rTg4510 mouse model, which underpins that in vivo monitoring of 4-repeat tau pathology in tau models is feasible with next generation tau radiotracers (5). From a biomechanical perspective, high individual tau-PET levels were linked with stronger atrophy in our cohort of PS19 mice, which highlights the link between tau and neurodegeneration. Furthermore, this finding shows that [^18^F]PI-2620 tau-PET signals in PS19 mice may still be underestimated since individual MRI based partial volume effect correction has yet to be established in mice.

One major goal of our investigation was to determine if tau-PET signals really derive from tau-positive cells. In this regard earlier tau-PET radiotracers such as [^18^F]THK5117 and [^18^F]AV1451 also showed increased signals in P301S or BiGT mice (20), but due to the uncovered off-target binding to monoamine oxidases (21), observed signal elevations could also be derived from activated immune cells, i.e. reactive astrocytes. As a major novelty, we applied cell sorting after in vivo tracer injection (22–24) to a tau radiotracer. Here we found that i) tau tracer binding was tremendously higher in neurons when compared to astrocytes, ii) neurons of PS19 mice had nearly 2-fold higher tracer uptake when compared to neurons of wild-type mice, and iii) single cell tracer uptake determined the in vivo tau-PET signal. Thus, scRadiotracing directly linked cellular binding to the translation towards an in vivo tau-PET signal. We note that our data cannot proof intra-neuronal binding to 4R-tau, but we deem neuronal off-target sources less likely when compared to immune cells (21), vessels, iron-associated regions, calcifications in the choroid plexus, or leptomeningeal melanin (18). Interestingly we also observed more tracer uptake of neurons when compared to astrocytes in wild-type mice, which could in magnitude indicate larger cell body volumes of neurons that subsequently contain higher levels of tau in physiological conditions as compared to astrocytes (25). scRadiotracing data in PS19 mice were in line with immunohistochemistry which also determined neurons as the major tau-positive cell type. Future work could directly correlate cellular radiotracer binding with the cellular amount of tau protein (22).

Our study includes the first sample of deceased patients with 4R-tauopathies and disease controls that allowed PET to autopsy correlation between in vivo [^18^F]PI-2620 signals with quantitative tau load ex vivo. Variable time intervals between PET imaging and death need to be considered as a limitation, since tau load could still change after PET imaging. Still, we obtained a strong correlation between in vivo PET signals and AT8 occupancy in the frontal cortex and the basal ganglia in our cohort consisting of six patients with PSP and two disease controls. In vivo tau-PET signals in patients without evidence of significant tau pathology ex vivo matched with the signal levels reported for healthy and disease controls (4). Noteworthy, one patient with definite PSP in autopsy was diagnosed as non-fluent primary progressive aphasia 3.2 years before death and one patient with definite PSP in autopsy was diagnosed as behavioral variant frontotemporal dementia 2.5 years before death. However, both also showed a PSP-like [^18^F]PI-2620 PET signal in the basal ganglia at the time of their diagnostic workup, which highlights the diagnostic value of PET imaging when 4R-tauopathies represent a possible differential diagnosis. Furthermore, [^18^F]PI-2620 PET also reliably captured AT8-positive tau (co)-pathology in the included TBK1 carrier.

To overcome the limitations of mixed pathology and variable intervals between PET and autopsy, we determined contributions of neuronal, oligodendroglial and astrocytic tau to human autoradiography signals, using a PSP dataset with very limited co-pathology. Previously, we already observed a higher agreement of post mortem neuronal tau covariance with tau-PET covariance when compared to astrocytic and oligodendroglial tau covariance in the same PSP samples (26). Building upon these results, we found a higher agreement between neuronal/oligodendroglial tau abundance and autoradiography signals when compared to astrocytic tau abundance in the frontal cortex. The most probable explanation of this finding was provided by similar analysis in the basal ganglia. Here, regions with neighboring large dense neurons even resulted in focal autoradiography signals whereas astrocyte or axonal dominated regions with AT8 of lower density revealed only low signal elevations. Similar effects were observed for oligodendroglia with higher AT8 density at the GM/WM boundary in contrast to lower intensity of AT8 for cortical astrocytes. Thus, faint processes of astrocytes and axon bundles likely suffer from partial volume effects, which hamper translation into a measurable tau radiotracer signal in vivo.

In particular, in cortical sections, we noticed pronounced autoradiography signals at the GM/WM boundary and we were able to correlate this finding to high proportions of AT8-positive oligodendroglia in this particular region. High oligodendroglial tau load in the white matter at the GM/WM boundary in 4R-tauopathies is also known from larger autopsy studies (16). In vivo transfer of this observation to [^18^F]PI-2620 tau-PET in patients with PSP showed the clinical relevance of this regional predominance, with higher effect sizes of PET signals extracted from the GM/WM boundary compared to the whole cortex regions (including grey matter and white matter). This should guide optimized frontal cortex target region selection of tau-PET analysis in 4R-tauopathies to increase PET sensitivity for cortical tau. In addition, dynamic imaging showed higher probability to detect clinically diagnosed 4R-tauopathies in vivo (4) compared to static imaging (27). This is also supported by lacking in vivo signals in PSP at later static imaging windows (28) and decreasing target signals with imaging time (29), supporting early (30) or dynamic scanning (4) to be preferred over late imaging windows. Thus, optimized target and reference tissues should be considered together with dynamic imaging to exploit the full diagnostic value of [^18^F]PI-2620 tau-PET in 4R-tauopathies.

In summary, we show that aggregated neuronal and oligodendroglial 4R-tau translates to a measurable tau-PET signal in patients with PSP and CBS, whereas astrocytic and axonal tau inclusions are a minor source of in vivo PET signals. Our novel approach of cell sorting after radiotracer injection can be readily used to test for the cell type specificity of novel radiotracers with 4R-tau affinity.

## Material and Methods

### Study design

In this translational study, we combined assessments of tau-PET, in vitro autoradiography, quantitative tau immunohistochemistry and cellular tracer uptake using a 4R-tauopathy mouse model and human samples consisting of patients with 4R-tauopathies and disease controls.

#### Small animal experiments

All small animal experiments had been approved by the local animal care committee of the Government of Upper Bavaria (Regierung Oberbayern, approval numbers: ROB-55.2-2532.Vet_02-15-210, ROB-55.2-2532.Vet_02-19-26). The experiments were overseen by a veterinarian and conducted in compliance with the ARRIVE guidelines and in accordance with the U.K. Animals (Scientific Procedures) Act, 1986 and associated guidelines, EU Directive 2010/63/EU for animal experiments. Animals were housed in a temperature- and humidity-controlled environment with 12h light-dark cycle, with free access to food (Ssniff Spezialdiäten GmbH, Soest, Germany) and water. Anesthesia was induced before [^18^F]PI-2620 application and maintained during the PET and MR scan with 1.5% isoflurane delivered via a mask at 3.5 L/min. All procedures were performed at the Department of Nuclear Medicine, Ludwig-Maximilian’s-University (LMU) Hospital, Munich. First, we conducted a longitudinal [^18^F]PI-2620 PET/MRI study in a 4R-tau mouse model (PS19) and age matched wild-type mice (n=10 each, all female), using regional tau-PET signals and volumetric measures as endpoints. PS19 transgenic mice feature the expression of mutated human microtubule-associated protein tau (MAPT) under the regulation of the mouse prion protein (Prnp) promoter. This transgene encompasses the P301S mutation linked to the disease and contains four microtubule-binding domains alongside an N-terminal insert (4R/1N) (13). Next, we performed immunohistochemistry in a subset of these PS19 and wild-type mice (n=4 each) to characterize regional tau abundance and cellular contributions to tau pathology. Cell sorting after tau radiotracer injection was applied in another subset of PS19 and wild-type mice (n=5 each) to determine the cellular origin of tau-PET signals.

#### Human analyses

A key experiment of the study consisted of a correlation analysis between regional [^18^F]PI-2620 tau-PET signals and abundance of fibrillary tau pathology in autopsy samples of patients with definite PSP (n=6) and disease controls (n=2, amyotrophic lateral sclerosis, TDP-43-positive frontotemporal lobe degeneration). In this sample, we performed a quantitative correlation analysis between tau-PET binding, autoradiography binding and abundance of AT8-positive tau pathology. An additional autopsy sample of deceased patients with PSP with limited co-pathology (n=16) was used to determine the contribution of tau-positive neurons and tau-positive astrocytes to [^18^F]PI-2620 autoradiography signals. To this end, abundance of AT8-positive tau pathology in subfields of frontal cortex and basal ganglia sections was differentiated between neurons and astrocytes by a data driven tissue classifier. Tissue samples of all autopsy cases investigated were provided by the Neurobiobank Munich, LMU Munich. They were collected according to the guidelines of the local ethical committee and usage of the material for this project was additionally approved (application number 19-244). Finally, the combined findings were used to evaluate the emerging grey matter/ white matter boundary target region for tau-PET assessment of 4R-tau pathology in the cortex of patients with PSP (n=17) in the contrast against controls (n=9). All patients and controls who received in vivo PET imaging provided informed written consent. The study was conducted in accordance with the principles of the Declaration of Helsinki, and approval was obtained from the local ethics committee (application numbers 17-569, 19-022).

### Small animal PET/MRI imaging

#### PET/MRI acquisition

All rodent PET procedures followed an established standardized protocol for radiochemistry, acquisition times and post-processing using a PET/MRI system. All mice were scanned with a 3T Mediso nanoScan PET/MR scanner (Mediso Ltd, Hungary) with a triple-mouse imaging chamber. Two 2-minute anatomical T1 MR scans were performed prior to tracer injection (head receive coil, matrix size 96 x 96 x 22, voxel size 0.24 x 0.24 x 0.80 mm³, repetition time 677 ms, echo time 28.56 ms, flip angle 90°). Injected dose was 12.7 ± 2.1 MBq for [^18^F]PI-2620 delivered in 200 µl saline via venous injection. PET emission was recorded in a dynamic 0-60 min window. Framing was 6×10, 2×30, 3×60, 5×120, 5×300, 5×600. List-mode data within 400-600 keV energy window were reconstructed using a 3D iterative algorithm (Tera-Tomo 3D, Mediso Ltd, Hungary) with the following parameters: matrix size 55 x 62 x 187 mm³, voxel size 0.3 x 0.3 x 0.3 mm³, 8 iterations, 6 subsets. Decay, random, and attenuation correction were applied. The T1 image was used to create a body-air material map for the attenuation correction. We longitudinally studied PS19 (n=10) and age-matched wild-type mice (n=10, WT; C57BL6) at 5.9, 7.7, 10.2 and 12.4 months of age. The sample size was selected based on the assumption of detecting a 10% difference between genotypes at the latest time-point with a power of 0.8, applying α of 0.05. No randomisation was used to allocate experimental units due to absence of any intervention. No dropouts were registered, hence all mice were able to be included for subsequent analysis.

#### PET/MRI analysis

Normalization of PET data was performed by calculation of volume of distribution (V_T_) images obtained from the full dynamic scan as described previously for different tracers (24, 31). In brief, we generated V_T_ images with an image derived input function using the methodology described by Logan et al. implemented in PMOD. The plasma curve was obtained from a standardized bilateral VOI placed in the left ventricle. A maximum error of 10% and a V_T_ threshold of 0 were selected for modelling of the full dynamic imaging data. Furthermore, 20-40 min static [^18^F]PI-2620 images were analyzed as a read-out matching the scRadiotracing normalization. We applied the striatum as a reference tissue to decrease the variability at the individual subject level and calculated V_T_ ratio as well as standardized uptake value ratio (SUVr) images. The reference tissue was validated by analyzation of V_T_ images in comparison of PS19 and WT mice, which confirmed no V_T_ differences between genotypes in the striatum (8.4 mm³). Predefined volumes-of-interest were delineated by spheres in the brainstem (4.2 mm³), and in the entorhinal cortex (2.8 mm³), guided by regions of the Mirrione atlas but eroded in order to avoid spill in of adjacent brain structures (**Supplemental Fig. 1A**). These target regions served for extraction of PET values for all mice.

MRI volumetric analysis was performed on coronal sections by manual delineation of the cerebellum, the brainstem and the striatum (each in three adjacent planes) using PMOD (**Supplemental Fig. 1B**). Cerebellum and brainstem were considered as a combined hindbrain region.

### scRadiotracing

#### Mouse brain dissociation

Five PS19 and five WT mice underwent scRadiotracing (22–24) immediately after the tau-PET scan. Adult Brain Dissociation Kit (mouse and rat) (Miltenyi Biotec, 130-107-677) was used for brain dissociation according to the manufacturer’s instructions. Adult mouse brains were dissected, briefly washed with Phosphate-buffered saline (PBS), cut into eight pieces, and dissociated with enzyme mix 1 and 2 using gentleMACS™ Octo Dissociator (Miltenyi Biotec, 130-096-427). The dissociated cell suspension was applied to pre-wet 100 µm Cell Strainer (Falcon, 352360). The cell pellet was resuspended using cold PBS and cold debris removal solution. Cold PBS was gently overlaid on the cell suspension. Centrifuged at 4°C and 3000×g for 10 minutes with acceleration and deceleration at 5. The two top phases were removed entirely. The cell pellets were collected and resuspended with 1 ml cold red blood cell removal solution followed by 10 minutes incubation. Cell pellets were collected for astrocyte and subsequent neuron isolation via magnetic activated cell sorting (MACS) (22–24).

#### Isolation of astrocytes

Adult Brain Dissociation Kit, mouse and rat (Miltenyi Biotec, 130-107-677) was used according to the manufacturer’s instructions. The prepared cell pellets were resuspended in 80 µl of AstroMACS separation buffer (Miltenyi Biotec, 130-117-336) per 10^7^ total cells. 10 μL of FcR blocking reagent were added and incubated for 10 minutes in the dark at 4°C. 10 μL of Anti-ACSA-2 MicroBeads were added and incubated for 15 minutes in the dark at 4°C. Cells were washed by adding 1 mL of AstroMACS separation buffer and centrifuged at 300×g for 5 minutes. Cell pellets were resuspended in 500 μL of AstroMACS separation buffer. The pre-wet MS columns (Miltenyi Biotec, 130-042-201) were placed at OctoMACS Separator (Miltenyi Biotec, 130-042-109). The cell suspensions were applied onto the column, followed by washing with 3 × 500 µL of AstroMACS separation buffer. The flow-through was collected containing non-astrocytic cells as an astrocyte-depleted fraction. The columns were removed from the magnetic field, and the astrocytes were flashed out using 3 ml of AstroMACS separation buffer.

#### Isolation of neurons

Neuron Isolation Kit, mouse (Miltenyi Biotec, 130-115-390) was used as previously reported (32), according to the manufacturer’s instructions. The astrocyte-depleted cell pellets were resuspended in 80 µl of PBS-0.5% Bovine Serum Albumin (BSA) buffer per 10^7^ total cells. 20 μL of Non-Neuronal Cells Biotin-Antibody Cocktail was added and incubated for 5 minutes in the dark at 4°C. Cells were washed and centrifuge at 300×g for 5 minutes. Cell pellets were again resuspended in 80 μL of PBS-0.5% BSA buffer per 10^7^ total cells. 20 μL of Anti-Biotin MicroBeads were added and incubated for 10 minutes in the dark at 4°C. The volume was adjusted to 500 µl per 107 total cells with PBS-0.5% BSA buffer and then proceed to magnetic separation. The pre-wet LS columns (Miltenyi Biotec, 130-042-401) were placed at QuadroMACS™ Separator (Miltenyi Biotec, 130-090-976). The cell suspensions were applied onto the columns. The columns were washed with 2 × 1 ml PBS-0.5% BSA buffer. The flow-through containing the unlabelled cells were collected as the neuron-enriched fractions. The columns were removed from the magnetic field, and the non-neuronal cells were flushed out using 3 ml of PBS-0.5% BSA buffer (24).

#### Gamma emission, flow cytometry and calculation of single cell tracer uptake

Radioactivity concentrations of cell pellets were measured in a high sensitive gamma counter (Hidex AMG Automatic Gamma Counter, Mainz Germany) relative to the activity in the whole brain, with decay-correction to time of tracer injection for final activity calculations.

Flow cytometry staining was performed at 4 °C. After gamma emission measurement, the cell suspension was centrifuged at 400*g* for 5 min and the supernatant was aspirated completely. The cell pellet was then resuspended in 100 µl of cold D-PBS containing fluorochrome-conjugated antibodies recognizing mouse CD11b and ACSA-2 (Miltenyi Biotec, 130-113-810 and 130-116-247) in a 1:100 dilution and incubated for 10 min at 4°C in the dark. Samples were washed with 2 ml of D-PBS and centrifuged for 5 min at 400*g*. Finally, cell pellets were resuspended in 500 μl of D-PBS and samples were immediately used for flow cytometry using a MACSQuant® Analyzer as quality control of MACS.

Measured radioactivity (Bq) of cell pellets was divided by the specific cell number in the pellet resulting in calculated radioactivity per cell. Radioactivity per cell was normalized by injected radioactivity and body weight (%ID*BW).

### Small animal immunohistochemistry

A total of four female PS19 mice and four wild-type mice were used for immunohistochemistry. 50 μm thick slices were cut in a sagittal plane using a vibratome (VT1200S, Leica Biosystems). Slices were treated with blocking solution (10% normal goat serum and 10% donkey serum in 0.3%Triton and PBS to a total volume of at least 200 μl per well/slice) for 3hrs at RT. The following primary antibodies were used: chicken anti-GFAP 1:500 (ab5541, Merck Millipore, Darmstadt, Germany), mouse anti-AT8 1:1000 (ab5541, Merck Millipore, Darmstadt, Germany), rabbit anti-MAP2 1:500 diluted in blocking solution (5% normal goat serum and 5% donkey serum in 0.3% Triton and PBS to a total volume of at least 200 μl per well/slice), applied to the slices and incubated for ∼48hrs at 4 °C on a horizontal shaker. Secondary antibodies, goat anti-rabbit Alexa Flour 488 (1:500), goat anti-chicken Alexa Fluor 555 (1:500), goat anti-mouse Alexa Fluor 647 (1:500) diluted in PBS, were applied. Slices were incubated for 2-3 hrs at RT on a horizontal shaker, protected from light. After 3 × 10 min washing with PBS, slices were mounted and cover slipped with fluorescence mounting medium containing DAPI (Dako, Santa Clara, USA).

Three-dimensional images were acquired with an Apotome microscope (Zeiss Oberkochen, Germany) using a 10× and 40× objective. The analysis programs Zeiss blue and ImageJ were used for quantification. Z-stack images (10 μm) were acquired with a 10× objective. Each signal (AT8 and GFAP) was quantified as the % area of the entire scanning frame.

Single optical sections were acquired using a 40x objective and the AT8 signal was analyzed as the perceptual are in MAP2 and GFAP-positive signal, respectively. To this end, we created a mask of the AT8 signal and transferred it into GFAP-positive astrocyte and MAP2 positive neuronal structures. After a local brightness/contrast adjustment and background subtraction we set a fixed threshold and calculated the AT8 area (%) inside the mask of the GFAP-positive astrocyte and MAP-positive neurons.

### Human Post-mortem samples

#### Human samples

For PET to autopsy analyses, we included all patients that received [^18^F]PI-2620 tau-PET prior to death, donation of the brain to the Munich brain bank and tissue workup by 31^st^ March 2024 (n=8; **Supplemental Table 1**). The formalin fixed and paraffin-embedded tissue blocks of the one hemisphere were used for AT8 and autoradiography analyses. Medial frontal gyrus and basal ganglia material including the globus pallidus was available for six patients with definite PSP, one patient with FTLD-TDP, and one patient with FTLD/MND-TDP. For in depth analyses of cellular and structural radiotracer signal origins, we selected samples with limited α-synuclein, TDP-43 and FUS pathology from the Munich brain bank. Limited β-amyloid pathology was tolerated, resulting in a total sample size of n=16 (**Supplemental Table 2**). Intact medial frontal gyrus material was available for fourteen patients and intact basal ganglia material including the globus pallidus was available for six patients.

#### Immunohistochemistry

Immunohistochemistry was performed on 4µm thick sections of formalin fixed and paraffin-embedded tissue using standard techniques. The immunohistochemical tau-staining was performed semi-automatically on a BenchMark device (Ventana, now Hoffmann-LaRoche, Basel, Switzerland) with mouse monoclonal AT8 antibody raised against hyperphosphorylated tau (Ser202/Thr205, 1:200, Invitrogen/Thermofisher, Carlsbad, CA, USA) on adjacent sections of those used in the ARG. The immunostained sections were digitized at 20x magnification with a Mirax Midi scanner (Zeiss, Carl Zeiss MicroImaging GmbH, Jena, Germany). For frontal cortex (medial frontal gyrus) and globus pallidus analyses, 8-12 regions of interest (subfields) were drawn manually per section and the AT8-positive tau load (%) was quantified using ZEN 3.4 blue edition software (Zeiss, Jena, Germany).

Similar to previous approaches, i.e. by Rittman et al (33), we aimed to subdivide AT8-positive tau load into different underlying cell types. Therefore we used semi-automatized object characterization and recognition to differentiate neurofibrillary tangles (NFT), coiled bodies (CB) and tufted astrocytes (TA) based on several parameters of morphological characteristics. Single NFT, CB and TA (n=15-20 objects per slice) were manually selected to define object thresholds including size (area), diameter, ellipse axis, perimeter, intensity, grade of circularity, roundness, and compactness. Due to substantial overlap of object characteristics, we subsequently defined NFT and CB as a combined group of AT8-positive cells with high density. We note that tau fragments (TF) were partially included in the TA channel, resulting in two analysis channels (NFT/CB and TA/TF). This was substantiated by correlation analysis between AT8-positivity and autoradiography signals in single subjects, that indicated similar associations of neuron and oligodendrocyte enriched regions with autoradiography signals. The final segmentation resulted in NFT/CB AT8-area-%, TA/TF AT8-area% and their intensities within 8-12 subfields per analyzed section.

For the correlation analysis between in vivo PET imaging and tau load in autopsy, composite regions of interest in the medial frontal gyrus and in the globus pallidus (internal and external part) were used.

#### Autoradiography

For direct comparison with the autoradiography signal, tau immunostaining from formalin fixed and paraffin-embedded tissue blocks from 16 PSP cases and three brain regions (frontal cortex, putamen, pallidum) was processed. For each patient and brain region, autoradiography with [^18^F]PI-2620 was performed on ≥4 sections as described previously (34). In brief, sections were incubated for 45 min (21.6 μCi/ml after dilution to a volume of 50 ml with phosphate buffered saline solution, pH 7.4, specific activity 480 ± 90 GBq/μmol), washed, dried, placed on imaging plates for 12 h and scanned at 25.0 µm resolution. Regions of interest were drawn on each sample using the AT8 staining of the adjacent section, thus serving for anatomical definition of subfields in the frontal cortex (grey matter and white matter). An AT8-negative region in the white matter was determined as reference region and ratios between subfield target regions and the reference region were calculated. Each subfield region was labeled as cortical or grey-matter/white matter boundary. Binding ratios were correlated with semiquantitative AT8 assessment using Pearson’s correlation after testing for normality and subject to a regression analysis (neuronal vs. astrocyte tau).

### Human PET imaging and analysis

#### Tau-PET acquisition and preprocessing

[^18^F]PI-2620 was synthesized as previously described (35). The injected dose ranged between 156 MBq and 223 MBq, applied as a slow (10 s) intravenous bolus injection. Positron emission tomography (PET) imaging was performed in a full dynamic setting (scan duration: 0-60 minutes post-injection) using a Siemens Biograph True point 64 PET/CT (Siemens, Erlangen, Germany) or a Siemens mCT (Siemens, Erlangen, Germany). The dynamic brain PET data were acquired in 3-dimensional list-mode over 60 minutes and reconstructed into a 336 x 336 × 109 matrix (voxel size: 1.02 x 1.02 x 2.03 mm^3^) using the built-in ordered subset expectation maximization (OSEM) algorithm with 4 iterations, 21 subsets and a 5 mm Gaussian filter on the Siemens Biograph and with 5 iterations, 24 subsets and a 5 mm Gaussian filter on the Siemens mCT. A low dose CT served for attenuation correction. Frame binning was standardized to 12×5 seconds, 6×10 seconds, 3×20 seconds, 7×60 seconds, 4×300 seconds and 3×600 seconds. Image derived input functions were generated by manual and automated extraction of the PET standardized uptake value (SUV) signal from the carotid artery over the 60 minute dynamic PET scan.

Via manual extraction, the blood activity concentration in the bilateral carotid artery was detected in early frames of the dynamic PET images, and spheres with a diameter of 5.0 mm were placed as volumes of interest (VOI) in the pars cervicalis of the internal carotid artery prior to entering the pars petrosal using PMOD version 4.2 (PMOD Technologies, Zürich, Switzerland). The activity concentration over time was calculated with the average values of the VOI.

#### Tau-PET quantification

Volume distribution (VT) images were calculated with the IDIFs using Logan Plots (36), which assume that the data become linear after an equilibration time t*. t* was fitted based on the maximum error criterion, which indicates the maximum relative error between the linear regression and the Logan-transformed measurements in the segment starting from t*. The maximum error was set to 10 %. The percent masked pixels were set to 0 %. The Putamen, which was defined by manual placement of a VOI (sphere with a diameter of 10 mm), served as tissue region.

All VT images were transformed to MNI space via the 20-40 min co-registration, using the established [^18^F]PI-2620 PET template (37). Automatized brain normalization settings in PMOD included nonlinear warping, 8 mm input smoothing, equal modality, 16 iterations, frequency cutoff 3, regularization 1.0, and no thresholding.

For PET to autopsy correlation analysis, VT ratio values (temporal white matter reference region (38)) were obtained in the medial frontal gyrus and in the globus pallidus as PSP target regions (see immunohistochemistry), predefined by the atlas of basal ganglia [11], the Brainnetome atlas [12], and the Hammers atlas [13]. The rationale was to employ a matched quantification strategy between PET and autoradiography.

#### GM/WM target region

All patients and controls used for this dedicated analysis underwent T1-weighted structural MRI on a 3T Siemens Magnetom PRISMA or SKYRA Scanner and [^18^F]PI-2620 tau-PET in a full dynamic setting (0–60 min post-injection) using pre-established standard PET scanning parameters (30). A detailed description of our in-house [^18^F]PI-2620 synthesis and quality control pipeline has been described previously (4, 39). Dynamic [^18^F]PI-2620 tau-PET imaging was acquired on a Siemens Biograph True point 64 PET/CT (Siemens, Erlangen, Germany) or a Siemens mCT scanner (Siemens, Erlangen, Germany) in 3D list-mode over 60 min together with a low dose CT for attenuation correction. From the dynamic PET image acquisition, we reconstructed late-phase tau-PET images for 20-40 min p.i. which were summarized into a single frame after motion correction (30, 40–42).

All images were screened for artifacts before preprocessing. T1-weighted structural MRI scans were bias-corrected, segmented into tissue types using the CAT12 toolbox (https://neuro-jena.github.io/cat12-help/). PET images were linearly co-registered to the corresponding T1 MRI data and intensity normalized using a pre-established inferior cerebellar reference region (26). Using T1-MRI data, surface reconstruction was performed using the CAT12 based cortical thickness pipeline. Surface reconstruction was performed for the GM/WM boundary, and then systematically shifted to the underlying white matter as well as to the GM/CSF boundary. Using these systematically shifted surfaces, we extracted PI-2620 tau-PET SUVRs for 200 regions of the cortical Schaefer atlas to systematically determine the tau-PET signal from the GM/CSF towards the GM/WM boundary and below.

### Statistics

Graph Pad Prism (V10, GraphPad Software, US) and SPSS (V27, IBM, US) were used for statistical analyses.

#### Mouse PET/MRI imaging

Mixed linear models (Graph Pad Prism) were used to test for age x genotype effects in PS19 and WT mice, including tau-PET binding and MRI volumes between 6 and 12 months of age as indices of interest. Pearson’s coefficient of correlation was calculated between brainstem tau-PET binding and brainstem volumes for all investigated time-points (separately for PS19 and WT mice), as well as between tau-PET binding and immunohistochemistry in n=4 PS19 mice. A sample size calculation was performed (G*Power) to determine the minimal age of detectability in tau-PET binding and MRI volume with cohorts of n=12 PS19 and n=12 WT mice (power 0.8, alpha 0.05; **Supplemental Fig. 2**).

#### Mouse immunohistochemistry

Regional coverage of AT8 and GFAP staining was compared between PS19 and WT mice by an unpaired Student’s t-test. Co-localization of AT8 with neurons (MAP2+) and astrocytes (GFAP+) was compared by an unpaired Student’s t-test.

#### Mouse scRadiotracing

Radiotracer uptake per neuron and astrocyte was compared between PS19 and WT mice using an unpaired Student’s t-test. Furthermore, within PS19 and WT mice radiotracer uptake was compared between neurons and astrocytes. Person’s coefficients of correlation were calculated between radioactivity per cell and tau-PET signals for the combined data of PS19 and WT mice as well as for the subset of PS19 mice. For whole-brain voxel-wise correlation analysis between cellular tracer uptake (n=5 PS19, n=5 WT), statistical parametric mapping (SPM) was performed using SPM12 routines (Wellcome Department of Cognitive Neurology, London, UK) implemented in MATLAB (version 2016). Individual SUVr images were subject to a linear regression analysis with cellular tracer uptake in neurons or astrocytes (%ID*BW) as a vector in the pooled cohort of PS19 and WT mice (Threshold: p < 0.005 uncorrected, k > 20 voxel). A paired t-test was used to compare PET radioactivity and extrapolated radioactivity of single cells in the cohort of five PS19 mice. PET radioactivity was determined as increases of counts per second (Bq) in PS19 compared to WT mice, Previously reported cell numbers of the mouse brain (43) were used to extrapolate the cellular radioactivity as a product of cellular abundance and radioactivity per cell.

#### Human PET to autopsy correlation

Pearson’s coefficients of correlation were calculated for the multimodal correlation between tau abundance in immunohistochemistry, autoradiography ratios and tau-PET signals.

#### Human autoradiography

Neuronal and astroglia tau abundance was compared with a paired t-test. Neuronal and astroglia tau abundance in subfields of the frontal cortex was correlated with autoradiography binding ratios in corresponding regions (total n=129; **Supplemental Fig. 3**). Additionally, a linear regression was performed with neuronal and astroglia tau abundance as predictors and autoradiography binding ratios as outcome variable. At the individual level, the correlation between neuronal and astroglia tau abundance and autoradiography binding ratios was performed in 8-12 subfields per subject. These individual correlations were analyzed as function of overall tau abundance (separately for neurons and astroglia). In basal ganglia regions, AT8 signal intensity and AT8 occupancy was correlated with autoradiography binding ratios.

#### Human PET target region

Differences in SUVR of n=5 different layers (GM/CSF boundary, GM towards CSF, GM towards WM, GM/WM boundary, below GM/WM boundary) were compared using paired t-tests.

### Data availability

All data needed to evaluate the conclusions of Fig. 1-8 are present in the paper and/or the Supplementary Materials. Imaging data will be shared in DICOM format upon reasonable request to the corresponding author.

## Author contributions

LS: performed autoradiography analyses and interpretation, designed autoradiography figure panels. JG: performed human PET data analyses and interpretation, designed PET figure panels. SH: participated in preclinical PET acquisition, image analysis and interpretation. LMB: performed scRadiotracing and analyzed scRadiotracing data, designed scRadiotracing figure panels. CK and LP: performed and interpreted small animal immunohistochemistry, designed small animal figure panels. AK, JK-P and LB processed and analyzed PET data. STK, LHK and AE: participated in preclinical PET acquisition. SK, CP, AB, AJ, AZ, FH, TG and JL performed clinical evaluation of all included patients and recruited patients for imaging and autopsy. GlBic and SS performed MRI acquisition and MRI data analyses. GBis, TvE, AD, OS and HB interpreted PET data and provided radiotracer supply. GR, LB, MW, JH and SR performed autopsy and definite diagnosis, provided and analyzed brain sections of patients, analyzed autoradiography. GUH, SR, NF and MB: conception and design, contributed to interpreting data, enhancing intellectual content of manuscript. All authors contributed with intellectual content and revised the manuscript. MB wrote the first draft of the manuscript with input of all co-authors.

## Acknowledgements

We thank Rosel Oos and Giovanna Palumbo for excellent technical support during small animal PET imaging. The authors thank the staff of the departments of nuclear medicine and neurology at the University Hospital LMU Munich. We thank the patients and their families.

