## Supplement for "Neuronal and oligodendroglial but not astroglial tau translates to in vivo tau-PET signals in primary tauopathies"

### Content

**Supplemental Figure 1 – Methodological details of small animal PET/MRI**

**Supplemental Figure 2 – Sample size estimation for assessment of tau burden and atrophy by small animal PET/MRI**

**Supplemental Figure 3 – Correlation between tau burden and tau-PET signals in PS19 mice**

**Supplemental Figure 4 – Subfield definition in autoradiography and AT8 sections**

**Supplemental Table 1 – Overview on samples of the PET to autopsy cohort**

**Supplemental Table 2 – Overview on samples of the autoradiography PSP cohort**

**Supplemental Figure 1**

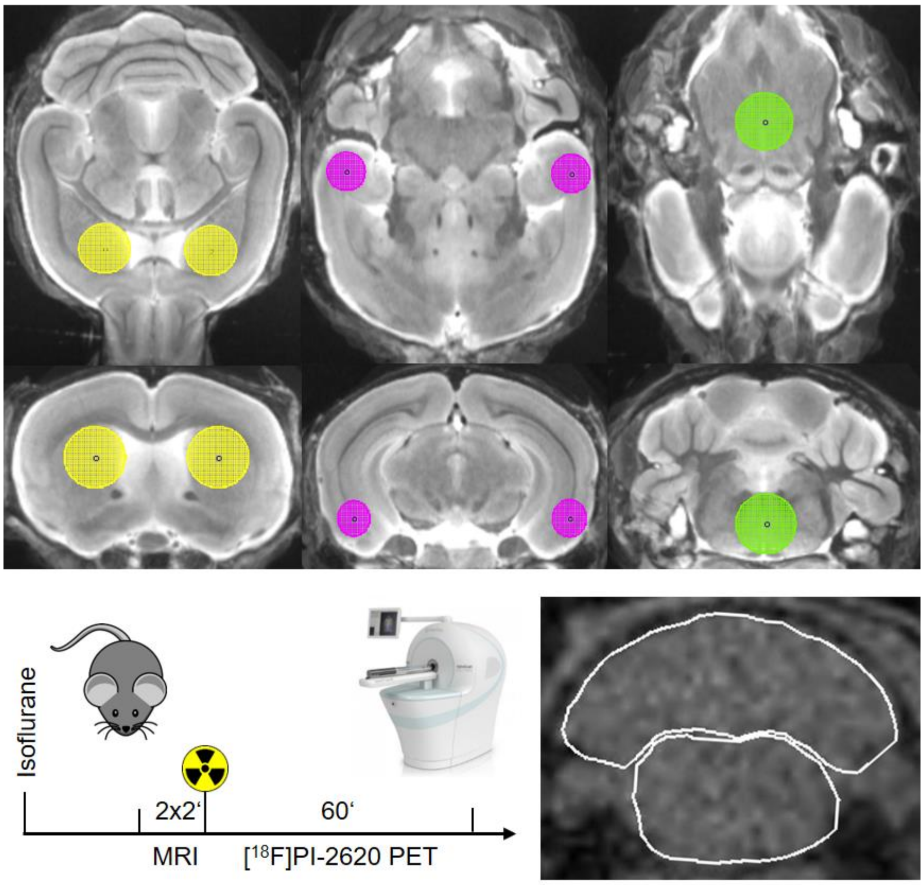


**Supplemental Figure 1 – Methodological details of small animal PET/MRI.** Upper panel shows definition of PET target (brainstem = green, entorhinal cortex = purple) and reference (striatum = yellow) regions. Lower panel shows timing of the PET/MRI acquisition protocol and delineation of brainstem and cerebellum volumes in a coronal plane.

### Supplemental Figure 2

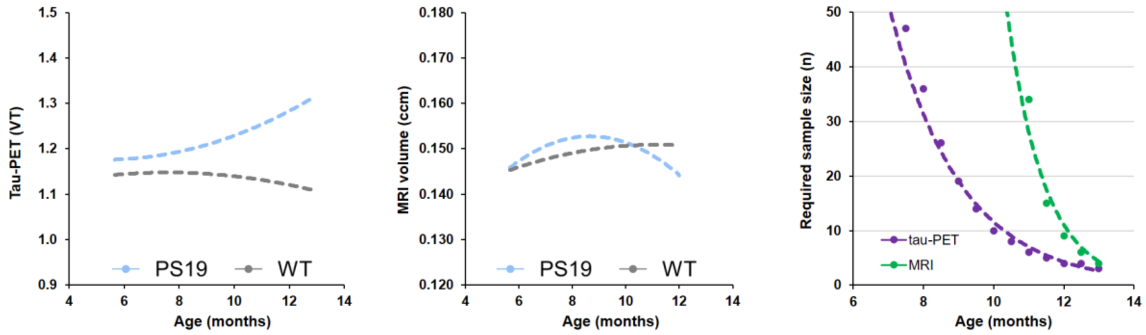


**Supplemental Figure 2 – Sample size estimation for assessment of tau burden and atrophy by small animal PET/MRI.** Trajectories of tau-PET signals and brain volumes as a function of age in PS19 and wild-type (WT) mice together with required sample size to detect significant differences in each modality at a power of 0.8 and alpha of 0.05.

### Supplemental Figure 3

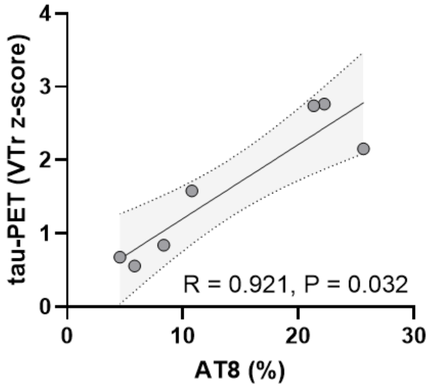


**Supplemental Figure 3 – Correlation between tau burden and tau-PET signals in PS19 mice.** Plot shows matched region pairs of AT8 area-% and [$^{18}$F]PI-2620 tau-PET change (volume of distribution ratio z-score) in frontal cortex (n=4) and hippocampus (n=3) of PS19 mice.

**Supplemental Figure 4**

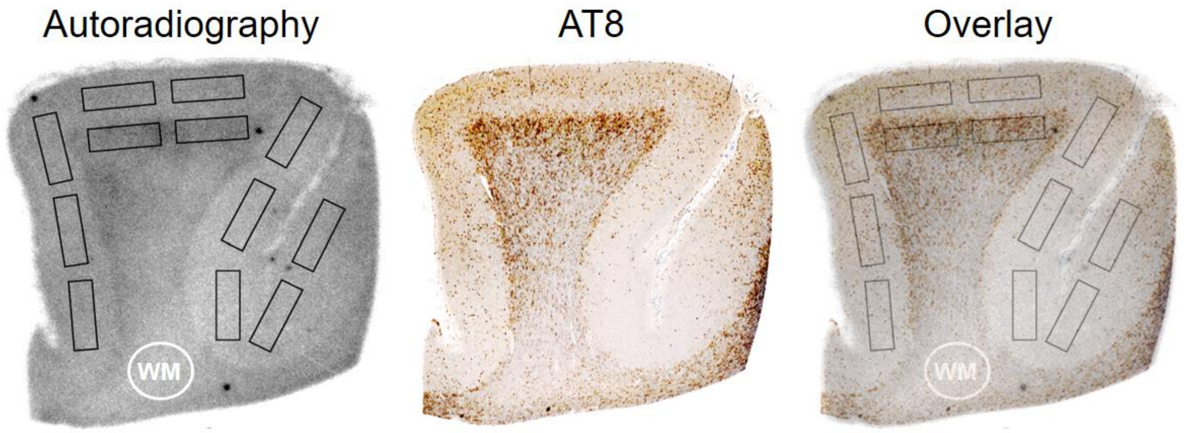


**Supplemental Figure 4 – Subfield definition in autoradiography and AT8 sections.** Example shows subfield definition in the grey matter and in the subcortical white matter of the medial frontal gyrus from a patient with definite PSP in autoradiography (left), the corresponding AT8 section (middle) and the overlay (right). Circular white matter (WM) region shows the reference region for determination of autoradiography target-to-white matter ratios. In this example both subfields in the autoradiography center were defined as grey matter/white matter boundary which showed high abundance of coiled bodies.

### Supplemental Table 1

| Case | Demographics | | | | | Diagnosis | | Autopsy determinants | | | Copathology | |
|---|---|---|---|---|---|---|---|---|---|---|---|---|
| | Gender | Age at PET (y) | Age at death (y) | Disease Duration (y) | Cause of death (death certificate) | Clinical diagnosis | Autopsy diagnosis | Brain weight (g) | Postmortem delay (h) | Fixation time (d) | Aβ/α-syn/TDP-43/FUS | Frontal cortex |
| #1 | female | 68 | 70 | 6 | Dysphagia | PSP-RS | PSP, CAA, AD (B&B 3), AGD | 1182 | 42 | 76 | | β-Amyloid |
| #2 | male | 64 | 66 | 4 | n.a. | PSP-RS | PSP, ARTAG, intranuclear inclusions of unclear etiology), discrete TDP-43 in the brainstem) | 1363 | 90 | 31 | | - |
| #3 | male | 70 | 71 | 6 | Atypical PD/ recurrent pneumonia | PSP | PSP | 1570 | 39 | 63 | | β-Amyloid |
| #4 | female | 73 | 77 | 6 | Cardiac arrhythmia | PSP-RS | PSP, AGD, ARTAG, subdural hemorrhage, AD (B&B 1) | 1170 | 25 | 71 | | β-Amyloid |
| #5 | male | 72 | 75 | 7 | Hearth failure, COVID-19 | bvFTD/nfvPPA | PSP | 1262 | 34-58 | 124 | | n.a. |
| #6 | male | 63 | 67 | 4 | n.a. | nfPPA | PSP; AD (B&B 3), mild AGD | 1512 | 30 | 48 | | - |
| #7 | female | 65 | 65 | 1 | ALS-FTD | ALS-FTD, PPA | FTLD/MND-TDP, AD (B&B 3), AGD, | 1261 | 48 | 88 | | - |
| #8 | female | 75 | 75 | 6 | Loss of food and fluids | bvFTD (TBK1-mutation) | - FTLD-TDP (Type A)<br>- Lewy body disease (Braak 5, neocortical)<br>-argyrophilic grain disease (AGD)<br>-ARTAG<br>-Central pontine myelinolysis<br>-Hippocampal sclerosis<br>-PART (B&B 2)<br>-multiple mikroiinfarcts | 1080 | 18 | 40 | | Synuclein, tau |

**Supplemental Table 1 – Overview on samples of the PET to autopsy cohort.** y = years, PD = Parkinson's disease, ALS = amyotrophic lateral sclerosis, FTD = frontotemporal dementia, PSP-RS = progressive supranuclear palsy Richardson syndrome, nf = non-fluent, PPA = primary progressive aphasia, bv = behavioral variant, AD = Alzheimer's disease, CAA = cerebral amyloid angiopathy, AGD = agyrophilic grain disease, ARTAG = aging-related tau astrogliopathy, B&B = Braak and Braak, TDP-43 = TAR DNA-binding protein 43, MND = motor neuron disease, PART = primary age-related tauopathy, Aβ = β-amyloid, APOE = apolipoprotein E, α-syn = alphasynuclein, n.a. = not available, „-„ = negative, co-pathology: FUS not examined

### Supplemental Table 2

| Case | Demographics | | | | Diagnosis | | Autopsy determinants | | | Copathology | | |
|---|---|---|---|---|---|---|---|---|---|---|---|---|
| | Gender | Age at death (y) | Disease Duration (y) | Cause of death | Clinical diagnosis | Autopsy diagnosis | Brain weight (g) | Postmortem delay (h) | Fixation time (d) | Aβ/α-syn/TDP-43/FUS | Frontal cortex | APOE |
| #1 | female | 67.3 | 7 | NA | PSP-RS | PSP, LBD Braak 4, CAA | 1290 | 14.27 | 646 | | Aβ: Vascular deposits | n/a |
| #2 | female | 70.9 | 5 | likely pulmonary embolism | CBS | PSP, AD alterations (Braak&Braak II, Thal 2, CERAD 0), mild arteriosclerosis | 1086 | 49.67 | 325 | | Aβ: some plaques | n/a |
| #3 | female | 74.9 | 10 | aspiration pneumonia | CBS | PSP, CAA | 1180 | 25.50 | 622 | | - | E3/E4 |
| #4 | male | 63.3 | 8 | NA | PSP-RS | PSP, AGD, ARTAG, arteriosclerosis, mild Aβ pathology (Thal 2) | NA | 18.75 | 518 | | Aβ: few diffuse plaques | n/a |
| #5 | male | 61.1 | 5 | NA | PSP-RS | PSP, old contusion frontal, macro-/microangiopathy | 1150 | 72.00 | 243 | | - | E3/E3 |
| #6 | female | 73.9 | 8 | NA | PSP-RS | PSP, AGD | 1140 | 55.00 | 375 | | - | E3/E3 |
| #7 | female | 87.4 | 13 | NA | PSP-CBS | PSP, LBD brain stem type, CAA | 1060 | 79.17 | 335 | | - | E2/E3 |
| #8 | female | 79.5 | 5 | cardiac arrest | PSP-RS | PSP, AGD | 1000 | 15.83 | 113 | | - | E3/E3 |
| #9 | male | 68.8 | 5 | cardiac arrest | FTLD, probably PSP | PSP, ischemic infarction CA1 left, arteriosclerosis | 1340 | 89.08 | 116 | | - | E3/E3 |
| #10 | male | 77.2 | 8 | bronchopneumonia, sepsis | PSP-RS | PSP, CAA, AD-alterations (Braak&Braak III, Thal 1), ARTAG, arterosclerosis, microinfarct left cerebellum | 1180 | 42.83 | 119 | | Aβ: few plaques | n/a |
| #11 | male | 75.7 | 8 | NA | FTD | PSP, AGD | NA | 97.52 | 246 | | - | n/a |
| #12 | male | 76.4 | 14 | aspiration pneumonia, subileus | PSP-RS | PSP | 1045 | 40.25 | 255 | | - | n/a |
| #13 | male | 75.8 | 5 | multiple organ failure | PSP-RS | PSP | 1440 | 40.82 | 477 | | - | n/a |
| #14 | male | 74.1 | 12 | NA | PSP-RS | PSP | 1200 | 11.58 | 675 | | - | n/a |
| #15 | male | 67.8 | 6 | cachexia | PSP-RS | PSP | 1460 | 37.50 | 195 | | - | n/a |
| #16 | male | 66.2 | 5 | cardiac and respiratory failure | PSP-RS | PSP | NA | 55.00 | 216 | | - | n/a |

**Supplemental Table 2 – Overview on samples of the autoradiography PSP cohort.** y = years, LBD = Lewy-body disease, ALS = amyotrophic lateral sclerosis, FTD = frontotemporal dementia, PSP-RS = progressive supranuclear palsy Richardson syndrome, nf = non-fluent, PPA = primary progressive aphasia, bv = behavioral variant, AD = Alzheimer’s disease, CAA = cerebral amyloid angiopathy, AGD = agyrophilic grain disease, ARTAG = aging-related tau astrogliopathy, B&B = Braak and Braak, TDP-43 = TAR DNA-binding protein 43, MND = motor neuron disease, PART = primary age-related tauopathy, Aβ = β-amyloid, APOE = apolipoprotein E, α-syn = alphasynuclein, NA = not available, „-„ = negative
